# Genetic Variants That Modify the Neuroendocrine Regulation of Foraging Behavior in *C. elegans*

**DOI:** 10.1101/2023.09.09.556976

**Authors:** Harksun Lee, Sonia A. Boor, Zoë A. Hilbert, Joshua D. Meisel, Jaeseok Park, Ye Wang, Ryan McKeown, Sylvia E. J. Fischer, Erik C. Andersen, Dennis H. Kim

## Abstract

The molecular mechanisms underlying diversity in animal behavior are not well understood. A major experimental challenge is determining the contribution of genetic variants that affect neuronal gene expression to differences in behavioral traits. The neuroendocrine TGF-beta ligand, DAF-7, regulates diverse behavioral responses of *Caenorhabditis elegans* to bacterial food and pathogens. The dynamic neuron-specific expression of *daf-7* is modulated by environmental and endogenous bacteria-derived cues. Here, we investigated natural variation in the expression of *daf-7* from the ASJ pair of chemosensory neurons and identified common variants in *gap-2*, encoding a GTPase-Activating Protein homologous to mammalian SynGAP proteins, which modify *daf-7* expression cell-non-autonomously and promote exploratory foraging behavior in a DAF-7-dependent manner. Our data connect natural variation in neuron-specific gene expression to differences in behavior and suggest that genetic variation in neuroendocrine signaling pathways mediating host-microbe interactions may give rise to diversity in animal behavior.

## Introduction

The molecular characterization of natural variants that affect behavioral traits provides a starting point to understand the cellular and organismal mechanisms driving differences in behavior (1–3). How variation in gene expression might be manifest in differences in behavior has been relatively unexplored in part because the relationship between neuronal gene expression and behavioral states is not well understood. Activity-dependent neuronal transcription has been shown to have pivotal roles in the development and plasticity of neural circuits, but less is known about how changes in gene expression play a role in shaping behavioral states (4). Moreover, whereas variation in levels of gene expression have been readily quantified on a genome-wide scale across evolutionarily diverse organisms in many tissues, including the nervous system, mechanistically bridging changes in levels of gene expression to differences in phenotypic traits has continued to be a major challenge (5–8).

Here, we focused on the characterization of natural variation in the neuron-specific expression levels of a single gene, *daf-7*, encoding at TGF-beta-related neuroendocrine ligand that regulates diverse aspects of *C. elegans* physiology (9–14). Neuronal expression of *daf-7* is dependent on multiple endogenous and environmental cues (9, 15–18). We have characterized the chemosensory signal transduction pathways involved in the regulation of dynamic *daf-7* expression in the ASJ neurons (15, 19). Recently, we characterized how the dynamic expression of DAF-7 in the ASJ neurons functions in a feedback loop that regulates the duration of exploratory roaming behavior in a two-state foraging behavior of *C. elegans* (20). In the present study, we sought to characterize natural variation in *daf-7* ASJ expression in order to test the hypothesis that changes in *daf-7* neuroendocrine gene expression could be a determinant of natural variation in behavior of *C. elegans*, with the anticipation that we might mechanistically connect the variants causing differences in *daf-7* expression to differences in DAF-7-dependent behavioral traits

## Results

### Natural variation in *daf-7* expression levels in the ASJ pair of neurons

We surveyed a set of wild strains of *C. elegans* for relative levels of *daf-7* expression in the ASJ neurons while feeding on the standard laboratory bacterial food *Escherichia coli* OP50. Previously, we found that for the laboratory wild-type strain N2 from Bristol, England, hermaphrodites do not exhibit detectable *daf-7* expression in ASJ neurons when feeding on *E. coli* OP50, but we have observed dynamic *daf-7* ASJ expression under different conditions, including the observation of *daf-7* ASJ expression in N2 hermaphrodites exposed to *P. aeruginosa* PA14 (15) and in N2 males feeding on *E. coli* OP50 (16). A genome-wide analysis of whole-animal gene expression levels across wild strains showed variation in *daf-7* expression (8). We observed a wide range of expression levels of *daf-7* in the ASJ neurons among wild strains sampled, with N2 exhibiting the lowest (undetectable) level of *daf-7* ASJ expression (Figs. 1a, b). Our survey was conducted using strains generated from a cross of an integrated transgenic multicopy fluorescent reporter under the control of the *daf-7* promoter (derived from the N2 strain) into each wild strain background. Our experimental approach thus precluded the observation of variation in *daf-7* expression due to local *cis-*acting loci because the reporter only had the regulatory regions of the N2 *daf-7* gene.

**Fig. 1.**
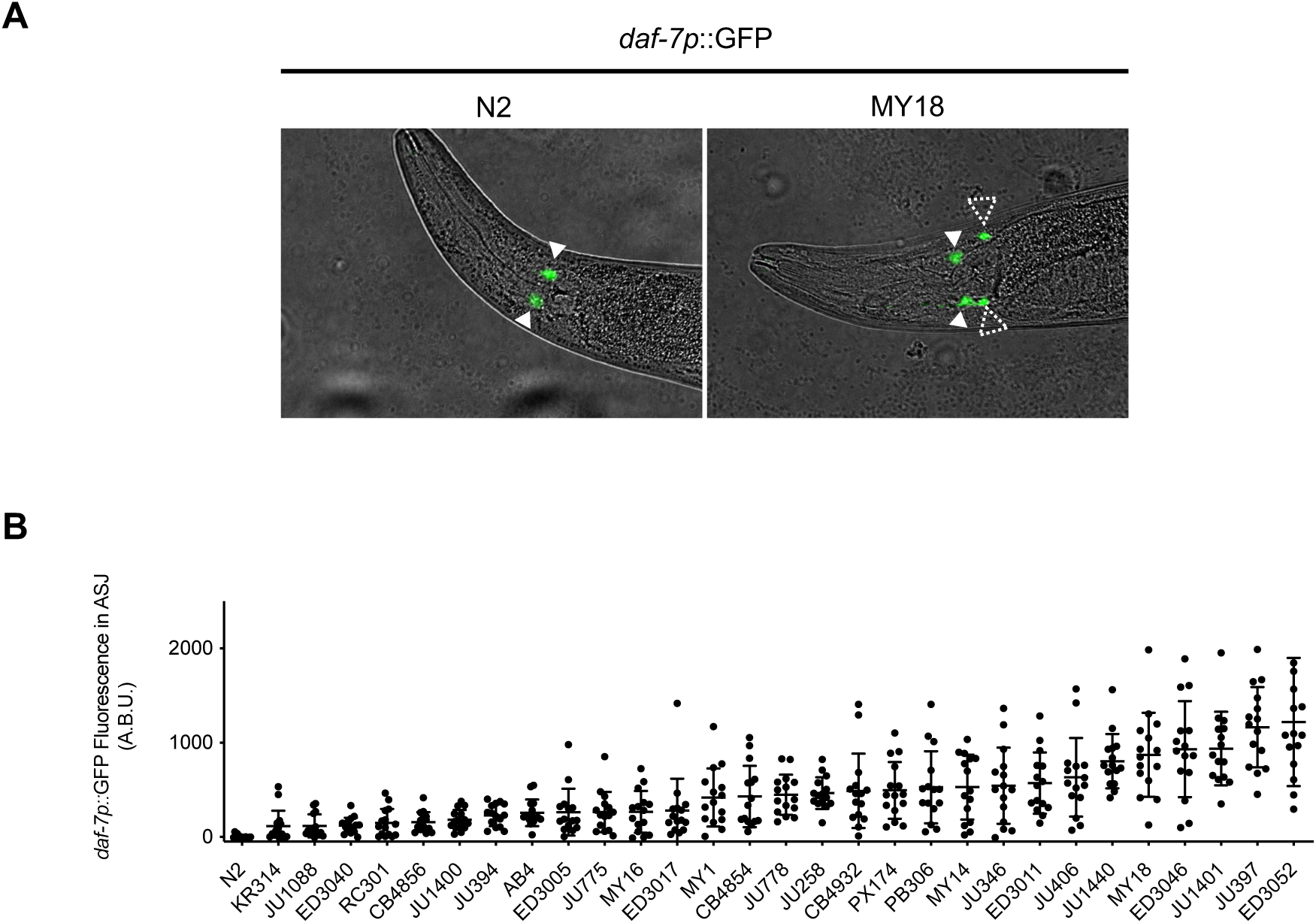
Natural variation in *daf-7* gene expression from the ASJ neurons in wild strains of *C. elegans*. (**A**) *daf-7p*::GFP expression pattern in N2 (left) and MY18 (right) genetic backgrounds. Filled triangles indicate the ASI neurons; dashed triangles indicate the ASJ neurons. Scale bar, 100μm. (**B**) Maximum fluorescence values of *daf-7p*::GFP in the ASJ neurons in the genetic background of indicated wild strains. Each dot represents an individual animal, and error bars indicate standard deviations.

### Identification of natural variants in *gap-2* modulating *daf-7* expression in the ASJ neurons

To define a molecular basis of natural variation in *daf-7* ASJ expression, we focused on differences in *daf-7* ASJ expression between N2 and MY18, a strain isolated from Münster, Germany. We generated 123 recombinant inbred lines (RILs) from multiple crosses between the N2- and MY18-derived strains (Fig. 2a) and quantified levels of *daf-7* ASJ expression in each RIL strain (Fig. 2b). RIL sequencing of MY18 was used to identify 5,716 variants that differed between the two parent strains (N2 and MY18), and a linkage mapping pointed to a broad quantitative trait locus (QTL) on chromosome X that contributed 11.5% of the observed differences in *daf-7* ASJ expression (Fig. 2c). In parallel, starting with an MY18-derived strain, we carried out multiple backcrosses with N2, selecting at each generation for the expression of *daf-7* ASJ expression, yielding near-isogenic lines (NILs) in the N2 genetic background defining a 447 kb interval on chromosome X: 9,513,008—X: 9,960,248 containing an MY18-derived locus that conferred increased *daf-7* ASJ expression in adult hermaphrodites feeding on *E. coli* OP50 (Fig. 2d).

**Fig. 2.**
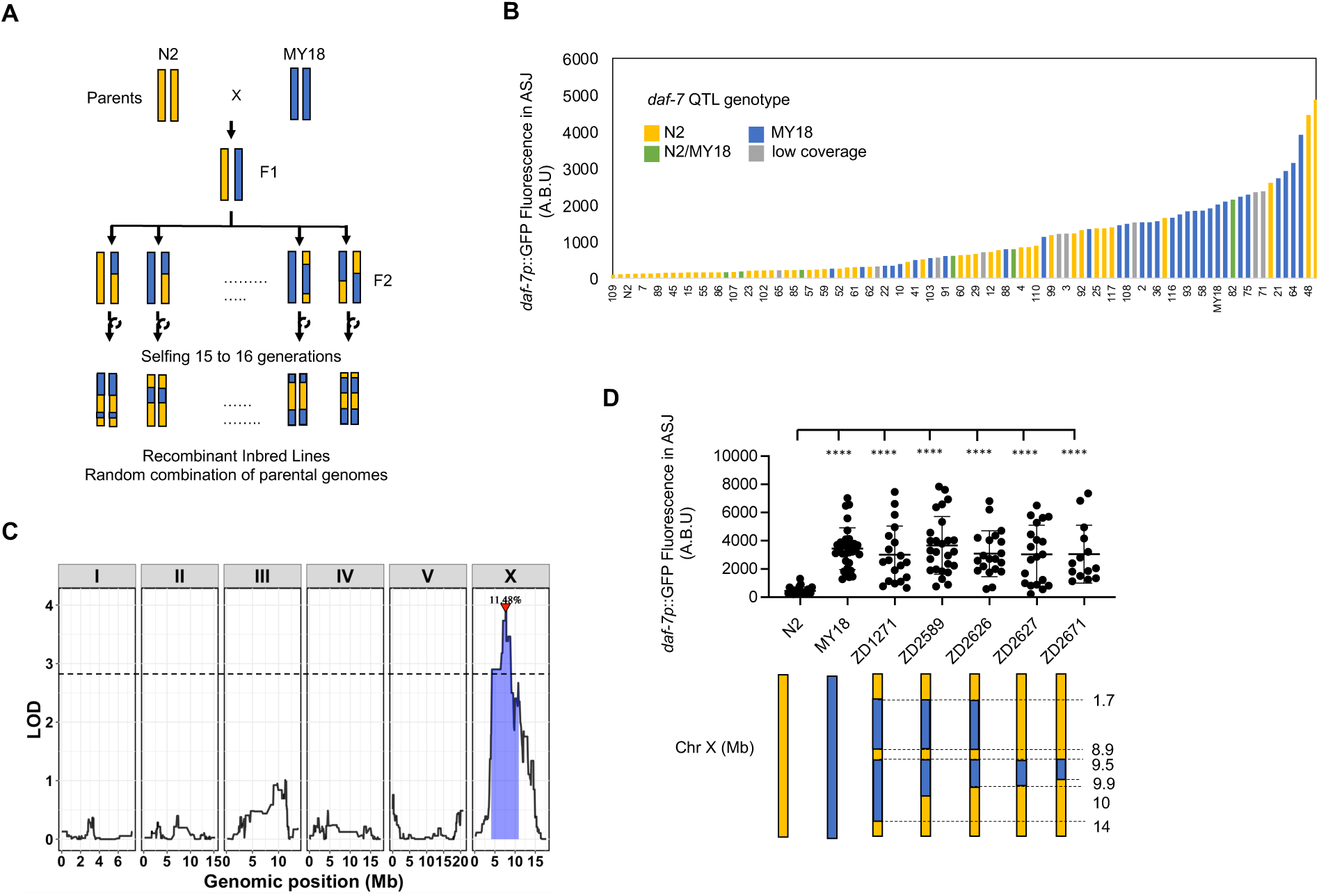
Identification of a genomic locus on chromosome X that affects *daf-7* expression in the ASJ neurons. (**A**) Diagram for constructing Recombinant Inbred Lines with strains derived from N2 and MY18 backgrounds. (**B**) Maximum fluorescence values *daf-7p*::GFP in the ASJ neurons of 123 recombinant inbred lines (RILs) and parental strains derived from N2 and MY18. The color of the bar represents the genotype at the QTL on chromosome X. (**C**) Linkage mapping results for *daf-7p*::GFP expression in the ASJ neurons. X-axis tick marks denote every 5 Mb. Significant QTL is denoted by a red triangle at the peak marker, and blue shading shows 95% confidence interval around the peak marker. The 5% genome-wide error rate LOD threshold is represented as a dashed horizontal line. LOD, log of the odds ratio. (**D**) Maximum fluorescence values *daf-7p*::GFP in the ASJ neurons of Near-Isogenic Lines (NILs). Each dot represents an individual animal, and error bars indicate standard deviations. ****p<0.0001 as determined by an unpaired two-tailed t-test compared to N2.

To identify the specific nucleotide differences contributing to the differential *daf-7* ASJ expression between the N2 and MY18 strains, we adopted a candidate variant approach, examining nucleotide differences predicted to cause coding changes in the 18 genes present in the narrowed interval on chromosome X. One of these genes was *gap-2*, encoding a conserved Ras GTPase-Activating-Protein (RasGAP) homologous to the SynGAP family. We identified a candidate variant present in MY18 that was predicted to affect an exon specific to two of the alternatively spliced isoforms of *gap-2*, *gap-2g* and *gap-2j*, and cause a putative substitution of a threonine in place of the serine at position 64 of GAP-2g and GAP-2j. First, we observed that a strain carrying the *gap-2(tm748)* deletion allele in the N2 background exhibited upregulated *daf-7* ASJ expression in adult hermaphrodite animals feeding on *E. coli* OP50 (Fig. 3g), leading us to evaluate the effect of the 64T variant in the N2 background. We observed that a strain carrying the *gap-2(syb4046)* allele, which had been engineered to carry the S64T change in the g/j isoforms of *gap-2* in the N2 genetic background, exhibited increased *daf-7* ASJ expression on *E. coli* OP50 (Fig. 3b, c). Conversely, when we examined the effect of engineering the 64S reciprocal change into the MY18 genetic background, we observed that *daf-7* ASJ expression decreased compared to what was observed in the MY18 strain (Fig. 3d). These data established that the S64T change in GAP-2g/j isoforms caused increased *daf-7* ASJ expression. We observed that the *gap-2(syb4046)* allele causes a dominant phenotype (Fig. 3f), where the *gap-2(tm748)* heterozygote conferred an intermediate level of *daf-7* ASJ expression compared to the wild-type or *gap-2(tm748)* homozygous strains (Fig. 3g).

**Fig. 3.**
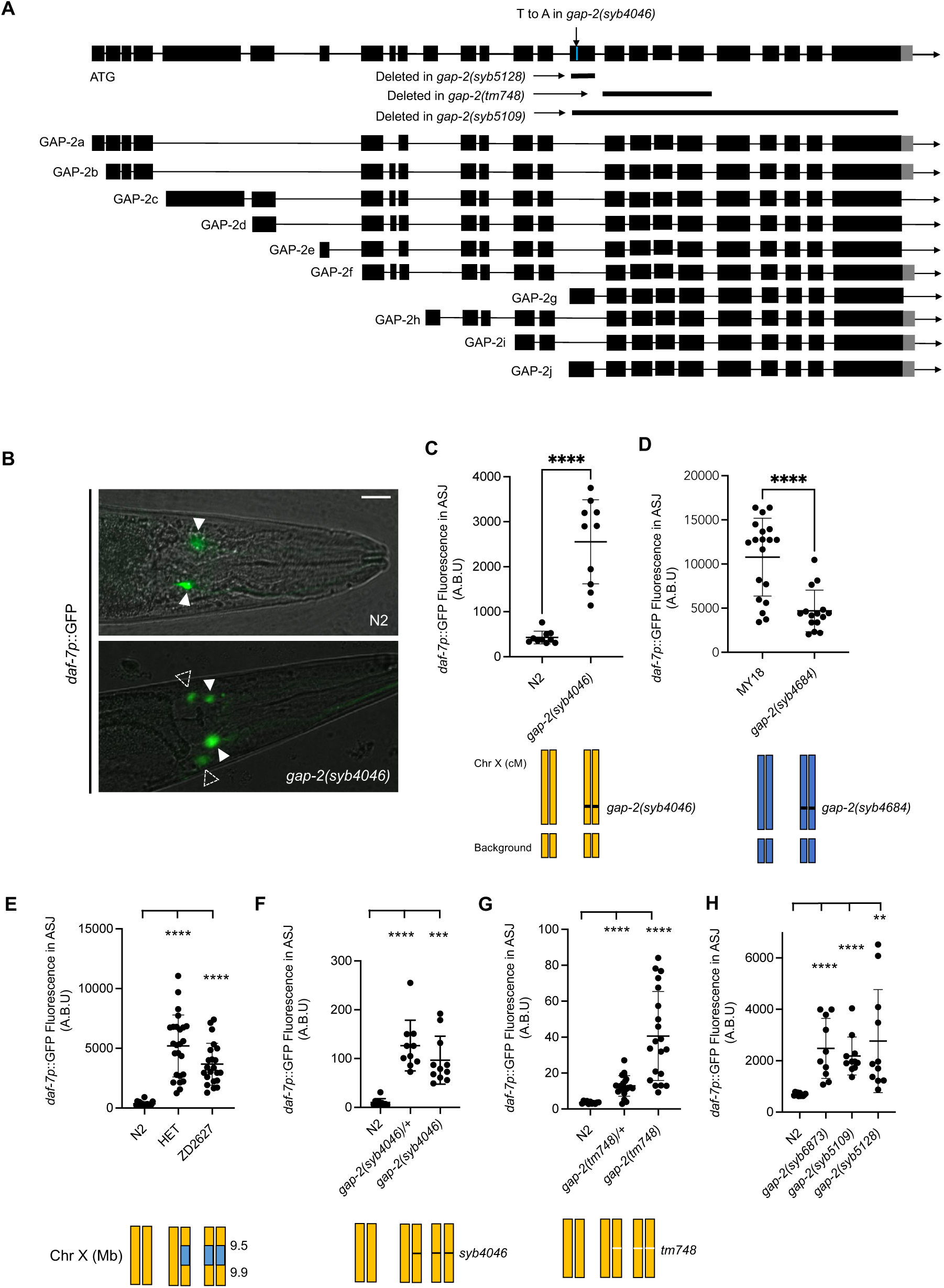
Natural variants in *gap-2* modulate *daf-7* expression in the ASJ neurons. (**A**) Genomic organization of multiple splice isoforms of *gap-2*. Deletion alleles of *gap-2* and the *syb4046* allele in the N2 background are depicted. (**B**) Representative images of *daf-7p*::GFP expression in N2 and *gap-2(syb4046)*. Filled triangles indicate the ASI neurons; dashed triangles indicate the ASJ neurons. Scale bar, 10 mm. (**C-H**) Maximum fluorescence values *daf-7p*::GFP in the ASJ neurons of indicated strains. HET is heterozygous of N2 and near isogenic line, ZD2627. Each dot represents an individual animal, and error bars indicate standard deviations. ****p<0.0001 and **p<0.01 as determined by an unpaired two-tailed t-test. Each genotype was compared to N2.

Based on the putative alteration of specifically *gap-2g* and *gap-2j* isoforms by the 64T variants, we examined whether other variants present in the affected exon specific to *gap-2g* and *gap-2j* isoforms might also influence *daf-7* ASJ expression. We found that 24 wild strains harbor a variant that causes a putative S11L change (Table 1b). To test the functional consequences of the S11L substitution in GAP-2, we generated the *gap-2(syb6873)* allele, in which the 11L variant was engineered in the N2 genetic background and observed that the 11L variant also conferred increased *daf-7* ASJ expression to a degree comparable to what was conferred by the 64T variant (Fig. 3h).

**Table 1a.** Strains that carry GAP-2(T64)

| Strains with T64 |
| --- |
| DL200, JU1213, JU1242, JU1400, JU1530, JU1666, JU1808, JU2001, JU2106, JU2131, JU2250, JU2478, JU2534, JU2566, JU2570, JU2575, JU2576, JU310, JU311, JU3128, JU3132, JU3140, JU3144, JU323, JU3795, JU440, JU792, JU847, MY16, MY18, MY2147, MY2453, MY2573, MY2741, MY772, MY795, NIC1, NIC1049, NIC1107, NIC166, NIC1698, NIC277, NIC3, NIC501, WN2064, WN2066, WN2086, WN2117 |

**Table 1b.** Strains that carry GAP-2(L11)

| Strains with L11 |
| --- |
| ECA369, JU258, JU2838, JU3166, JU3167, JU3169, MY1, NIC2021, NIC2076, NIC252, NIC266, NIC267, NIC268, NIC269, NIC274, NIC276, QG2813, QG2832, QG2837, QG2841, QG2846, QG4228, WN2063, XZ1735 |

### *gap-2* natural variants cause differences in a foraging behavioral trait

Having established that two *gap-2* variants promoted *daf-7* ASJ expression, we sought to connect these variants that alter gene expression to corresponding *daf-7*-dependent behavioral traits. *C. elegans* exhibits a two-state foraging behavior that alternates between an exploratory roaming state and an exploitative dwelling state on bacterial food (21, 22). A natural variant in *exp-1* (23) and a laboratory-derived mutation in *npr-1* (24) have been previously shown to modify the proportion of time that animals spend in the roaming and dwelling states. *daf-7* mutants have been shown to spend an increased proportion of time in the dwelling state compared to the N2 wild-type strain (25, 26). Recently, we have shown that the expression of *daf-7* in the ASJ neurons is part of a dynamic gene expression feedback loop that promotes increased duration of the roaming state (20).

We observed that the MY18 strain exhibited a marked increase in the proportion of time spent in the roaming state relative to the N2 strain (Fig. 4a, d and h). The ancestral 215F allele of *npr-1* is known to contribute to increased roaming behavior when compared to the N2 strain that carries a laboratory-derived 215V mutation in *npr-1* (24, 27, 28). To examine the effect of the 64T *gap-2* variant on foraging behavior, we observed the strain carrying the *gap-2(syb4046)* S64T allele in the N2 genetic background and found this strain spent an increased proportion of time in the roaming state relative to the N2 strain (Fig. 4a, b and h). In addition, we observed that the strain carrying the *gap-2(syb4684)* T64S allele in the MY18 genetic background spent a decreased proportion of time in the roaming state (Fig. 4d, e and h). These data establish that the 64T variant in *gap-2* promotes an increase in the proportion of time that the animal is in the roaming state relative to the dwelling state when compared with the 64S allele of *gap-2* in the N2 or MY18 genetic backgrounds. We also observed that the strain carrying the *gap-2* allele with the S11L variant, *gap-2(syb6873)*, in the N2 genetic background also increased the proportion of time that animals spent in the roaming state, to a degree comparable to what was observed for the 64T variant in the N2 background (Fig. 4a, c and h). These data identify two distinct variants acting on the same exon specific to *gap-2g* and *gap-2j* isoforms having comparable effects on *daf-7* ASJ expression and foraging behavior.

**Fig. 4.**
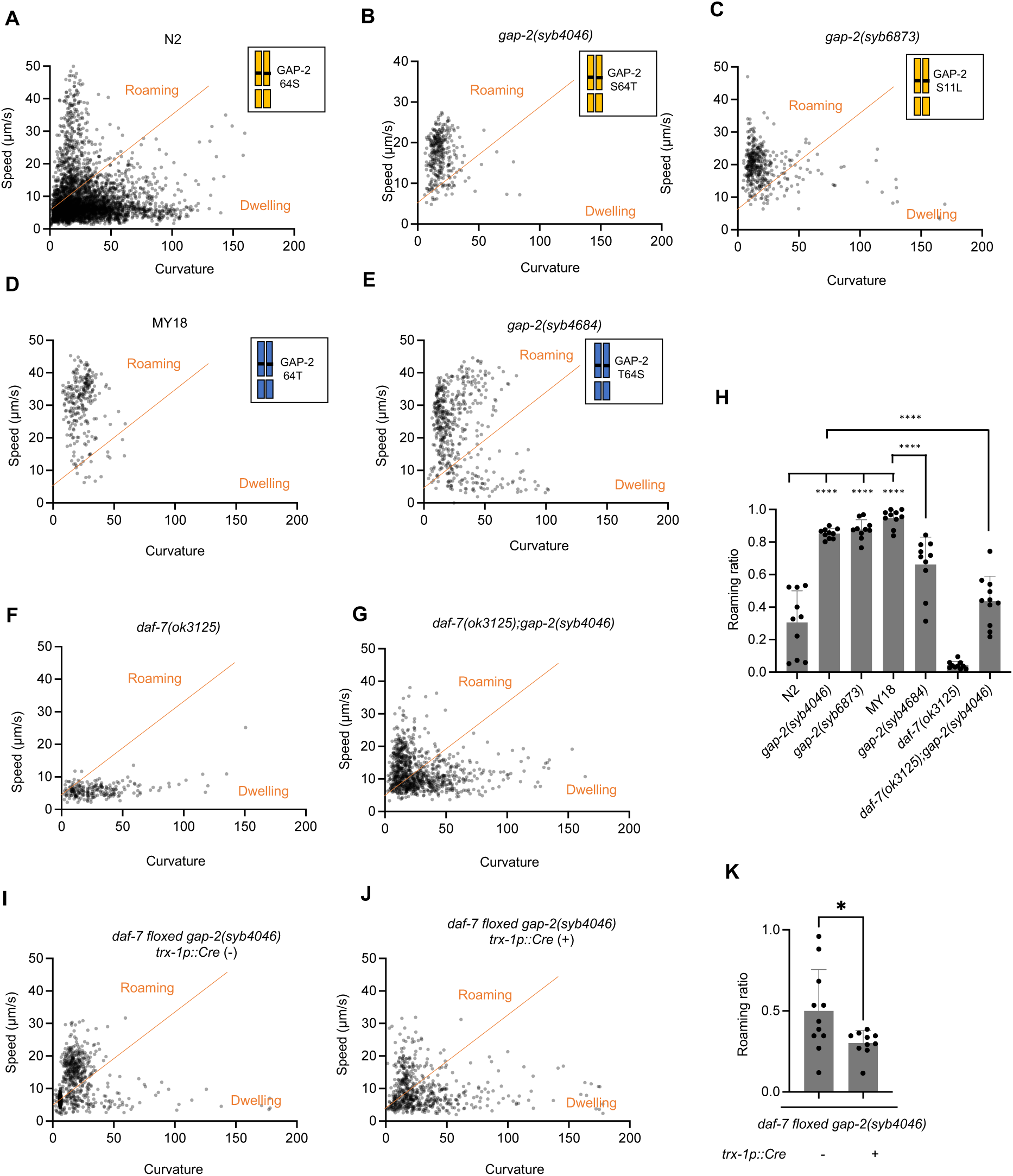
*gap-2* variants affect foraging behavior. (**A-G, I, J**) Scatter plot of speed and curvature of each indicated strain. Each dot represents average speed and body curvature during 10 seconds. The orange line was determined on total N2 data and used for analyses of all the strains. Chromosome diagrams indicate *gap-2* allele and genetic background (Yellow: N2, Blue: MY18). (**H**) Roaming ratio of N2, *gap-2(syb4046), gap-2(syb6873),* MY18, *gap-2(syb4684)*, *daf-7(ok3125)* and *daf-7(ok3125); gap-2(syb4046)*. Each dot represents average roaming ratio of one animal. ****p<0.0001 as determined by an unpaired two-tailed t-test. (**K**) Roaming ratio of *daf-7* floxed *gap-2(syb4046)* with or without *Ptrx-1::Cre* transgene expression. Each dot represents average roaming ratio of one animal. *p<0.05 as determined by an unpaired two-tailed t-test.

### *gap-2* variants cause differences in foraging behavior by modifying *daf-7* neuroendocrine gene expression

To determine whether the *gap-2* variants modified roaming behavior by their effects on *daf-7* ASJ expression, we first examined the roaming and dwelling behavior of the strain carrying the T64 *gap-2* variant and a loss-of-function *daf-7* allele in the N2 genetic background. We observed that this *gap-2(syb4046); daf-7(ok3125)* double mutant exhibited diminished roaming behavior compared with a strain carrying only the *gap-2(syb4046)* allele in the N2 background (Fig. 4f, g and h). Next, we examined the roaming behavior of a strain carrying the *gap-2(syb4046)* allele and a floxed *daf-7* locus, with and without a transgene expressing Cre under an ASJ-specific (*trx-1p*) promoter. We observed a diminished proportion of roaming behavior in the animals carrying a selective knockout of *daf-7* from the ASJ neurons (Fig. 4i, j and k). These data suggest that the effect of the *gap-2(syb4046)* variant on *daf-7* ASJ expression contributes to the foraging behavior trait difference between the N2 and MY18 strains, although the partial suppression of the increased roaming conferred by the *gap-2(syb4046)* variant suggests that the variant might also act through DAF-7-independent mechanisms to promote roaming behavior.

### *gap-2* variants act cell-non-autonomously to modulate neuroendocrine gene expression and foraging behavior

To determine the site-of-action where *gap-2* variants modify neuroendocrine gene expression and foraging behavior, we first defined the expression pattern of *gap-2* by GFP-tagging endogenous GAP-2 at its C-terminus. We observed expression in multiple cells of the nervous system and vulval cells (Supplemental movie 1a, b), consistent with prior reports of the tissue expression pattern of *gap-2* (29). To determine the cells in which the *g-* and *j-* isoforms of *gap-2* were expressed, we inserted a stop codon in the exon preceding the first exon of the *g-* and *j-* isoforms, which was expected to cause only translation of GFP-tagged G- and J- isoforms of GAP-2 (Fig. 5a, b). This strain exhibited GFP expression in a restricted set of cells, including presumptive expression in the ADE neuron pair based on cell location and morphology. We confirmed expression of the putative G- and J- isoforms of GFP-tagged GAP-2 in the ADE neuron pair through colocalization with the expression of the *dat-1p*::*mCherry* transgene that is expressed in the ADE neuron pair (Fig. 5c).

**Fig. 5.**
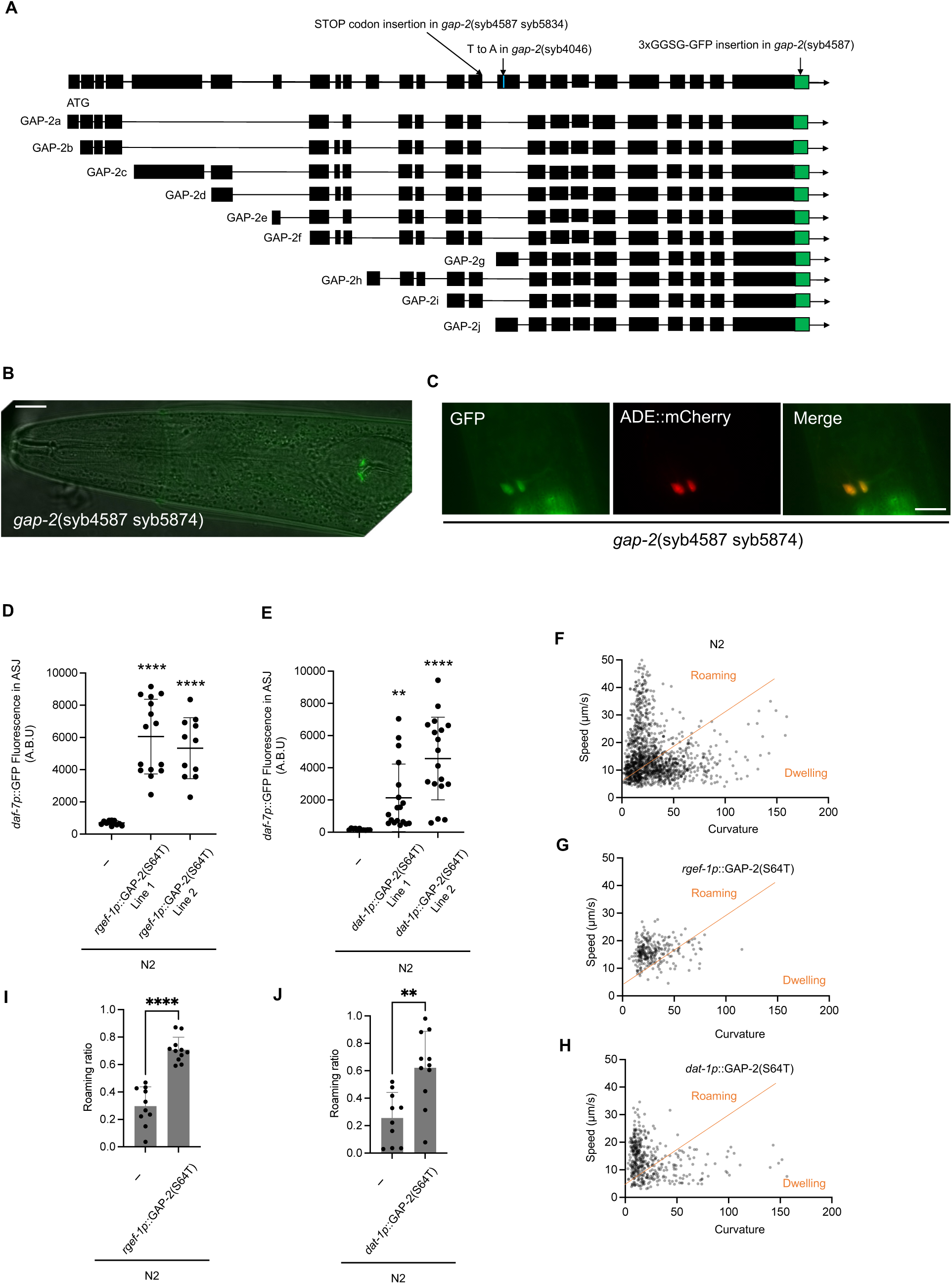
Natural variants in *gap-2* act in the ADE pair of neurons was sufficient to modulate *daf-7* ASJ expression and foraging behavior. (**A**) Diagram of the *gap-2* genomic locus with C- terminal GFP tagging and stop codon insertion engineered to facilitate the determination of expression pattern. (**B**) GFP tagged GAP-2g/j isoform expression of the *gap-2(syb4587 syb5834)* strain. Scale bar, 10 mm. (**C**) Colocalization of GAP-2g/j isoform and *dat-1p*::mCherry reporter. Scale bar, 10 mm. (**D**) Maximum fluorescence values *daf-7p*::GFP in the ASJ neurons of pan- neuronal GAP-2(64T) expressing transgenic strains. Each dot represents an individual animal, and error bars indicate standard deviations. ****p<0.0001 as determined by an unpaired two- tailed t-test. Each genotype was compared to WT. (**E**) Maximum fluorescence values *daf- 7p*::GFP in the ASJ neurons of ADE neuronal GAP-2(64T) expressing transgenic strains. Each dot represents an individual animal, and error bars indicate standard deviations. ****p<0.0001 and **p<0.01 as determined by an unpaired two-tailed t-test. Each genotype was compared to WT. (**F-H**) Scatter plot of speed and curvature of each indicated strain. Each dot represents average speed and body curvature during 10 seconds. Orange line has been determined on total N2 data and used for analysis of all the strains. (**I**) Roaming ratios of pan- neuronal GAP- 2(S64T) expressing transgenic strains. Each dot represents an individual animal, and error bars indicate standard deviations. ****p<0.0001 as determined by an unpaired two-tailed t-test. (**J**) Roaming ratios of ADE neuronal GAP-2(S64T) expressing transgenic strain. Each dot represents an individual animal, and error bars indicate standard deviations. ****p<0.0001 as determined by an unpaired two-tailed t-test.

We used the expression pattern information to determine the neurons in which expression of the T64 variant of GAP-2 was sufficient to confer increased *daf-7* ASJ expression and roaming behavior. Because the 64T variant causes a dominant effect on *daf-7* expression, we used a transgene expressing the *gap-2 g*-isoform cDNA carrying the 64T variant under the control of the pan-neuronal *rgef-1* promoter in the N2 genetic background. We also expressed the *gap-2 g*- isoform cDNA carrying the 64T variant under the control of the ADE neuronal *dat-1* promoter in the N2 background. We observed that the expression of the 64T GAP-2 sequence under the control of both pan-neuronal and ADE neuronal promoters, in the N2 genetic background, conferred *daf-*7 ASJ expression (Fig 5d, e) and an increased proportion of time spent in the roaming state (Fig. 5i, j). These data suggested that expression of the GAP-2 variant in the ADE neurons was sufficient to act cell-non-autonomously to alter the expression of *daf-7* from the ASJ neurons and to modify foraging behavior.

### Prevalence and geographical distribution of *gap-2* variants

The 64T *gap-2* variant is present in 48 out of 550 isotype reference strains in the *Caenorhabditis* Natural Diversity Resource (30), whereas the 11L *gap-2* variant is found in 24 of these strains. We also identified two additional rare variants, ENN39E and P19S, which are also predicted to affect the *g* and *j* isoforms of *gap-2*. We examined the geographic origins of these 72 wild strains and observed enrichment for the T64 allele in Africa and Europe (Supplementary Fig S1a). We found that the 11L and ENN39E variants have also been observed in divergent strains from the Hawaiian Islands. Because *daf-7* expression is affected by the bacterial diet, we investigated enrichment in specific natural substrates and found that the T64 allele more often found in strains isolated from compost as compared strains isolated from rotting fruit or leaf litter (Supplementary Fig S1b).

## Discussion

Our data illustrate how changes in neuronal gene expression caused by natural variants underlie the mechanism by which the variants can exert differential effects on behavioral traits. In particular, we show that natural variants cause differences in behavior by cell-non-autonomously modulating expression levels of a neuroendocrine regulatory ligand in two neurons of *C. elegans*. The systematic analysis of gene expression as a quantitative trait has been generally conducted on a genome-wide scale enabled by RNA-seq-based methodology (32). Our focus on the expression of a single gene enabled us to use a transgenic fluorescent reporter to examine levels of neuron-specific expression in live animals and facilitated subsequent mapping and molecular characterization. Moreover, *daf-7* expression has been previously demonstrated to exhibit dynamic neuron-specific expression and have key functional roles in diverse physiological processes (5, 8–18). Roles for neuromodulators in shaping circuits that govern behavior have been implicated from genetic studies, and our data suggest that variants causing differences in expression levels of genes that encode neuromodulators represent candidate variants that cause differences in behavior.

GAP-2 is one of three GTPase-Activating Proteins in the *C. elegans* genome that stimulate the LET-60 Ras GTPase to reduce EGF growth factor signaling (29, 33). GAP-2 is orthologous to Disabled 2 interactive protein (DAB2IP), a member of the SynGAP family of GAPs, which has been implicated in a range of disease states (34, 35). The similar dominant effects on *daf-7* ASJ expression for both of the 64T and 11L natural variants as compared to the recessive effects of a deletion of *gap-2* suggest that 64T and 11L variants confer reduced GAP-2 activity. We speculate that the 64T and 11L GAP-2G and/or GAP-2J isoforms act as dominant-negative proteins that might bind to LET-60 but not activate the GTPase and compete with wild-type GAP-2. Both *let-60(n1046)* gain-of-function and *let-60(n2021)* reduction-of-function alleles exhibit pronounced locomotory phenotypes in the presence of bacterial food (36), precluding the analysis of genetic interactions between *gap-2* and *let-60*, but the presence of locomotory defects is consistent with *gap-2* affecting Ras signaling in the ADE neurons to alter foraging behavior.

Our data also illustrate how the presence of multiple alternatively spliced transcripts of *gap-2* might facilitate the emergence of genetic variants that exert effects in a restricted set of neurons. Whereas GAP-2 appears to be widely expressed in the nervous system and in other tissues, the GAP-2g and/or GAP-2j isoforms exhibit restricted expression in the ADE pair of sensory neurons, and two variants, both 64T and 11L, affect an exon specific to these two isoforms.

The neuron-specific expression of *daf-7* is modulated by multiple bacteria-derived cues, including environmental and internal food cues (20) and secondary metabolites produced by pathogenic bacteria (15), and DAF-7 signaling contributes to exploratory behaviors, including lawn avoidance (15), mate-searching (16), and roaming behavior (20). The genetic variants in *gap-2* that we have identified that modulate *daf-7* expression from the ASJ neurons and promote roaming behavior are found in many strains throughout the *C. elegans* species, suggesting that the variants may confer a fitness advantage in diverse bacterial environments. The enrichment of these *gap-2* variants in strains isolated from compost relative to strains isolated from rotting fruit or leaf litter substrates could be because *C. elegans* isolated from compost are more commonly found as dauer larvae (37). We speculate that animals carrying *gap-2* variants might gain advantage in compost environments where increased exploratory behavior might be beneficial in acquiring nutrients. Our data suggest that diversity in behavioral traits associated with host interactions with microbes (38) can emerge from variants affecting the regulation of neuronal gene expression that is modulated by bacteria-derived food and pathogen cues.

## Acknowledgments

We thank Bob Horvitz and the Caenorhabditis Genetics Center, which is funded by the NIH Office of Research Infrastructure Programs (P40 OD010440), for strains. We thank the *Caenorhabditis* Natural Diversity Resource, which is funded by the NSF Capacity grant (2224885), and WormBase for critical genomic and natural variation data. We thank members of the Kim lab for discussions.

## Author contributions

HL: Conceptualization, Formal analysis, Validation, Investigation, Visualization, Methodology, Writing—original draft, Writing—review and editing SAB, ZAH, JDM, JP, SEJF: Investigation, Visualization, Writing – review and editing YW: Validation, Investigation, Methodology, Visualization ECA: Supervision, Visualization, Methodology, Writing—review and editing DHK: Conceptualization, Supervision, Funding acquisition, Visualization, Methodology, Writing—original draft, Writing—review and editing

## Data and materials availability

All data are available in the main text or the supplementary materials.

Strains that have A instead of T in the locus of Chromosome X:9,513,008 (Table 1a) and strains that have T instead of C in the locus of Chromosome X:9,512,850 (Table 1b). Database search from CeNDR variant annotation (30).

## Supplementary Figure

**Fig. S1.**
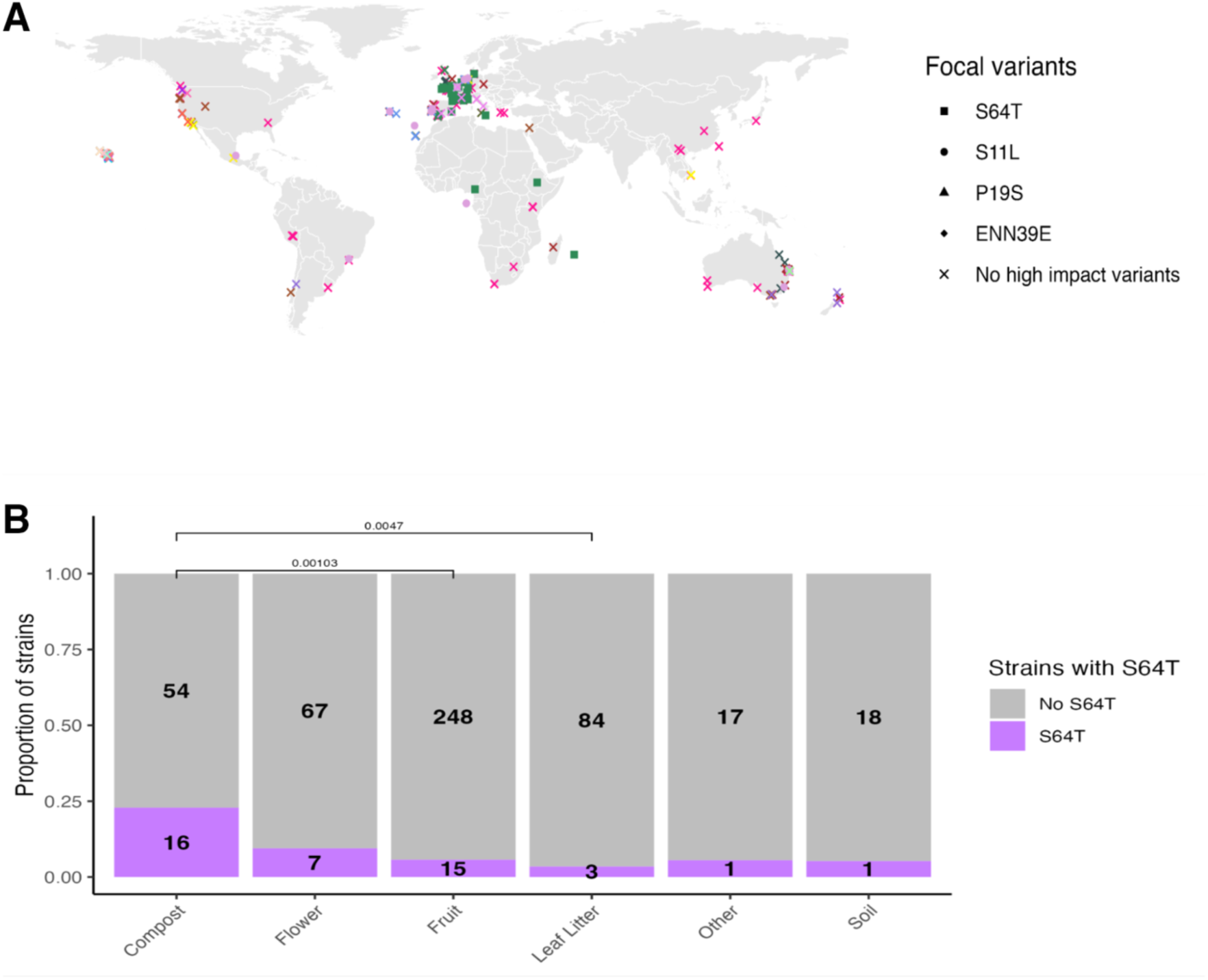
Environmental enrichment of *gap-2* S64T allele is observed in natural populations. (**A**) *gap-2* haplotypes of wild *C. elegans* strains are represented by colored shapes (32 unique haplotypes across 550 strains) and are plotted by their sampling location. Shapes distinguish the presence or absence of a naturally occurring variant predicted to affect the splicing or protein coding sequence of *gap-2* g/j. (**B**) Substrate enrichment of the S64T across wild strains. A Fisher’s exact test was used to evaluate significance.

**Supplemental movie 1a**

https://drive.google.com/file/d/11KbckYKS6j_-6UNXZb2oPG-mMeeZZgnL/view?usp=sharing

**Supplemental movie 1b**

https://drive.google.com/file/d/11KbckYKS6j_-6UNXZb2oPG-mMeeZZgnL/view?usp=sharing

## Supplementary Materials and Methods

### C. elegans culture

All strains were grown at 20°C on nematode growth medium plates seeded with Escherichia coli OP50 bacteria. OP50 was inoculated into 50ml of B broth and grown 24 hours at 37°C. See Supplemental Table 1. for a complete list of strains used in this study.

### *daf-7p*::GFP expressing wild isolate strain construction

To generate *daf-7p*::GFP carrying natural isolate reporter strains, each wild strain of interest was initially crossed to FK181 to introduce the *ksIs2* transgene. Progeny carrying the transgene were then backcrossed to the parental wild strain 8 times.

### Near Isogenic Lines construction

ZD1271 was made by backcrossing the ZD841 8 times to FK181 by following the *daf-7p*::GFP expression in ASJ phenotype.

ZD2589 was made by crossing the ZD1271 with FK181 then screened the strains for having N2 genotype region in between the Chr. X 9.5Mb∼14Mb.

ZD2626 was made by crossing ZD2589 with FK181 then screened the strains for having N2 genotype in between Chr. X 9.5Mb∼ 12Mb.

ZD2627 was made by crossing ZD2589 with FK181 then screened the strains for having N2 genotype in Chr. X 1.7Mb∼ 8.9Mb.

ZD2671 was made by crossing ZD2589 with FK181 then screened the strains for having N2 genotype in between Chr. X 9.5Mb∼ 10Mb.

### Linkage mapping

*daf-7p*::GFP expression in RILs were measured and whole genome sequencing result of each RIL strains were used for linkage mapping. A total of 123 RILs were examined for the level of *daf-7p*::GFP expression in ASJ neurons as described above. Linkage mapping was performed for *daf-7p*::GFP expression in ASJ neurons using R package linkage mapping (www.github.com/AndersenLab/linkagemapping<u>) as</u> previously described (39). QTL were detected using the fsearch function. This function calculates the logarithm of the odds (LOD) scores for each genetic marker and each trait as -n(ln(1-R2)/2 ln (10)) where R is the Pearson correlation coefficient between the RIL genotypes at the marker and trait values (40). A significant threshold based on a 5% genome-wide error rate was calculated. QTL were identified as the marker with the highest LOD score above the significance threshold. This marker was then integrated into the model as a cofactor and mapping was repeated until no significant QTL were detected. The annotate lods function was used to calculate the effect size of each QTL. 95% confidence intervals were defined by a 1.5-LOD drop from the peak marker.

### *daf-7p*::GFP expression imaging

For quantification of *daf-7* expression in animals, day-one young adult animals were mounted and anesthetized in levamisole. The animals were imaged at 40x using a Zeiss Axio Imager Z1 microscope. 15-20 animals were imaged for each condition or strain. For quantification, maximum intensity values of GFP within the ASJ neurons were calculated using FIJI software (<u>Schindelin et al., 2012</u>); Fiji, RRID:<u>SCR_002285</u>).

### Roaming/dwelling assay

Day-one adults were placed on 10 cm NGM plates, on which fresh OP50 had been uniformly spread the day before the experiment. These plates contained a 6 cm copper ring to keep the animals within the field of view for recording, The worms were transferred to inside of the ring on the assay plates 40 minutes before to move to worm tracker to avoid the initial reaction of worms to picking and transfer. On average 10 worms were recorded. We recorded the region inside the copper ring at 3.75 frames per second for 90 minutes. Videos were analyzed using MBF Biosciences WormLab software. Speed and mid body bending angle were averaged over 10 second intervals. Values for each 10 second interval were plotted on a scatter plot of speed (y axis) and bending angle (x axis). Quantification of fraction of time spent roaming was done by the diagonal line whose placement was based on the distribution of points of the distribution of N2. Equation for this line is y=0.3*x+5. Points falling above the line were classified as roaming and those points below the line were classified as dwelling.

### Statistics

All statistical analysis was performed using the Graphpad Prism software. Statistical tests used for each experiment are listed in the figure legend.

**Supplemental Table 1.** Complete list of *C. elegans* strains used in this study.

| Strain Name | Description | Source |
| --- | --- | --- |
| N2 | Wild type | Caenorhabditis Genetics Center (CGC) |
| FK181 | *ksIs2[pdaf-7::gfp;rol-6(su1006)]* | CGC |
| ZD1190 | KR314 crossed FK181 for 8 times to introduce *ksIs2* | This study |
| ZD838 | JU1088 crossed FK181 for 8 times to introduce *ksIs2* | This study |
| ZD1097 | ED3040 crossed FK181 for 8 times to introduce *ksIs2* | This study |
| ZD1075 | RC301 crossed FK181 for 8 times to introduce *ksIs2* | This study |
| ZD710 | CB4856 crossed FK181 for 8 times to introduce *ksIs2* | This study |
| ZD1635 | JU1400 crossed FK181 for 8 times to introduce *ksIs2* | This study |
| ZD1028 | JU394 crossed FK181 for 8 times to introduce *ksIs2* | This study |
| ZD837 | AB4 crossed FK181 for 8 times to introduce *ksIs2* | This study |
| ZD1120 | ED3005 crossed FK181 for 8 times to introduce *ksIs2* | This study |
| ZD834 | JU775 crossed FK181 for 8 times to introduce *ksIs2* | This study |
| ZD840 | MY16 crossed FK181 for 8 times to introduce *ksIs2* | This study |
| ZD1073 | ED3017 crossed FK181 for 8 times to introduce *ksIs2* | This study |
| ZD1187 | MY1 crossed FK181 for 8 times to introduce *ksIs2* | This study |
| ZD1119 | CB4854 crossed FK181 for 8 times to introduce *ksIs2* | This study |
| ZD839 | JU778 crossed FK181 for 8 times to introduce *ksIs2* | This study |
| ZD1096 | JU258 crossed FK181 for 8 times to introduce *ksIs2* | This study |
| ZD1638 | CB4932 crossed FK181 for 8 times to introduce *ksIs2* | This study |
| ZD1188 | PX174 crossed FK181 for 8 times to introduce *ksIs2* | This study |
| ZD1027 | PB306 crossed FK181 for 8 times to introduce *ksIs2* | This study |
| ZD1189 | MY14 crossed FK181 for 8 times to introduce *ksIs2* | This study |
| ZD1026 | JU346 crossed FK181 for 8 times to introduce *ksIs2* | This study |
| ZD1025 | ED3011 crossed FK181 for 8 times to introduce *ksIs2* | This study |
| ZD1636 | JU406 crossed FK181 for 8 times to introduce *ksIs2* | This study |
| ZD1074 | JU1440 crossed FK181 for 8 times to introduce *ksIs2* | This study |
| ZD841 | MY18 crossed FK181 for 8 times to introduce *ksIs2* | This study |
| ZD1633 | ED3046 crossed FK181 for 8 times to introduce *ksIs2* | This study |
| ZD1186 | JU1401 crossed FK181 for 8 times to introduce *ksIs2* | This study |
| ZD1632 | JU397 crossed FK181 for 8 times to introduce *ksIs2* | This study |
| ZD1637 | ED3052 crossed FK181 for 8 times to introduce *ksIs2* | This study |
| ZD1271 | Near Isogenic Line (MY18 region Chr. X 1.7~8.9Mb and Chr. X 9.5Mb~14Mb) | This study |
| ZD2589 | Near Isogenic Line (MY18 region Chr. X 1.7~8.9Mb and Chr. X 9.5Mb~ 12Mb) | This study |
| ZD2626 | Near Isogenic Line (MY18 region Chr. X 1.7~8.9Mb and Chr. X 9.5~+10Mb) | This study |
| ZD2627 | Near Isogenic Line (MY18 region Chr. X 9.5Mb~ 10Mb) | This study |
| ZD2671 | Near Isogenic Line (MY18 region Chr. X 9.5Mb~ 9.9Mb) | This study |
| ZD2683 | *ksIs2;gap-2(syb4046)* | This study/SunyBiotech |
| PHX4046 | *gap-2(syb4046)* | This study/SunyBiotech |
| PHX5109 | *gap-2(syb5109)* | This study/SunyBiotech |
| PHX5128 | *gap-2(syb5128)* | This study/SunyBiotech |
| PHX6873 | *gap-2(syb6873)* | This study/SunyBiotech |
| PHX4684 | *gap-2(syb4684)* | This study/SunyBiotech |
| PHX4587 | *gap-2(syb4587)* | This study/SunyBiotech |
| PHX5834 | *gap-2(syb4587 syb5834)* | This study/SunyBiotech |
| JN147 | *gap-2(tm748)* | National BioResource Project (NBRP) |
| RB2302 | *daf-7(ok3125)* | CGC |
| ZD2687 | *ksIs2;gap-2(tm748)* | This study |
| ZD2704 | N2;Ex[*rgef-1p::GAP-2(S64T)* + PQZ22] | This study |
| ZD2705 | N2;Ex[*ceh-2p::GAP-2(S64T)* + PQZ22] | This study |
| ZD2706 | N2;Ex[*trx-1p::GAP-2(S64T)* + PQZ22] | This study |
| ZD2707 | FK181;Ex[*rgef-1p::GAP-2(S64T)* + PQZ22] | This study |
| ZD2708 | FK181;Ex[*ceh-2p::GAP-2(S64T)* + PQZ22] | This study |
| ZD2709 | FK181;Ex[*trx-1p::GAP-2(S64T)* + PQZ22] | This study |
| ZD2713 | N2;EX[*dat-1p::GAP-2(S64T)* + PQZ22] TG line 1 | This study |
| ZD2714 | N2;EX[*dat-1p::GAP-2(S64T)* + PQZ22] TG line 2 | This study |
| ZD2715 | N2;EX[*dat-1p::GAP-2(S64T)* + PQZ22] TG line 3 | This study |
| ZD2716 | FK181;Ex[*dat-1p::GAP-2(S64T)* + PQZ22] TG line 1 | This study |
| ZD2717 | FK181;Ex[*dat-1p::GAP-2(S64T)* + PQZ22] TG line 2 | This study |

### Sequence information of strains generated by CRISPR

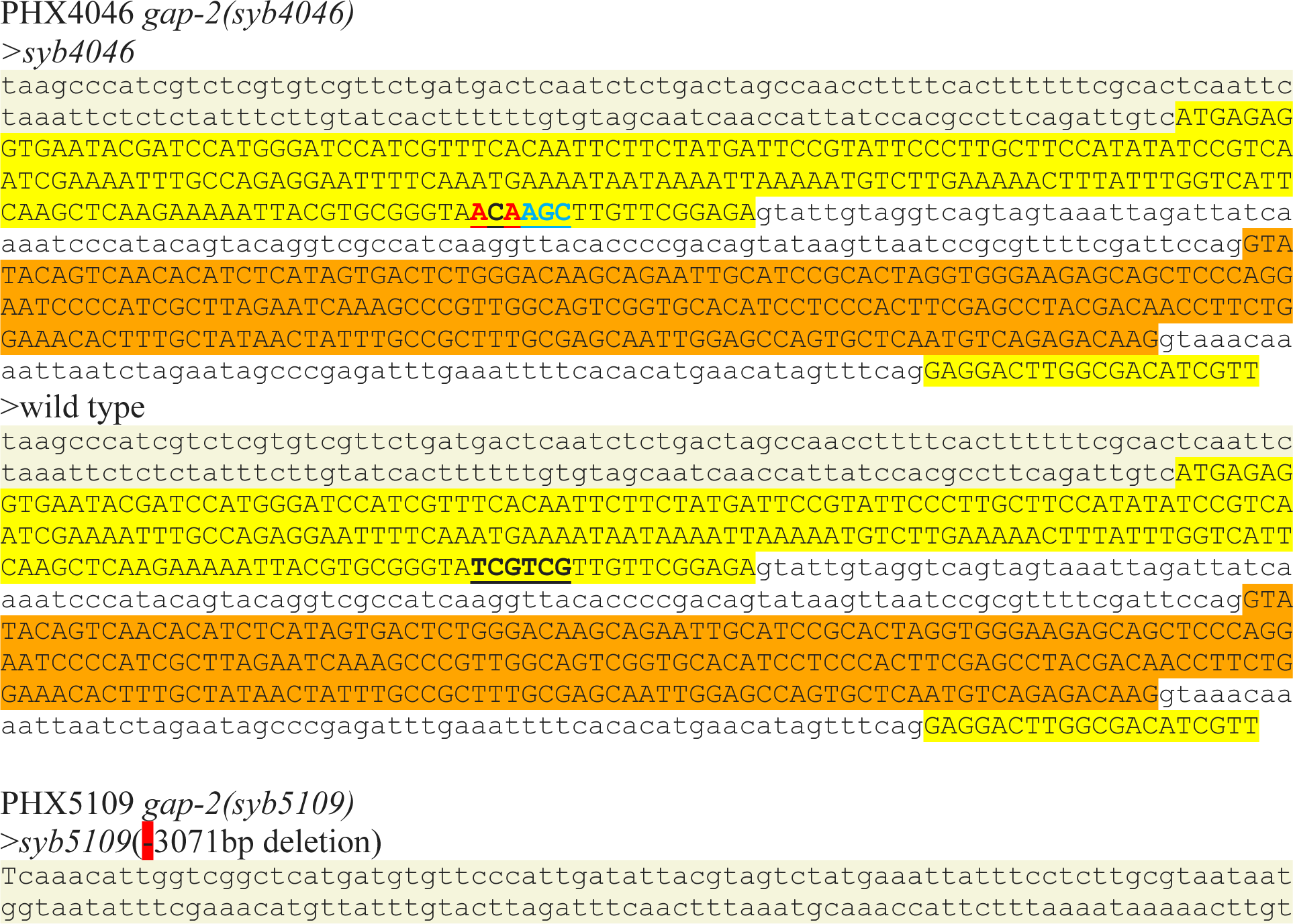
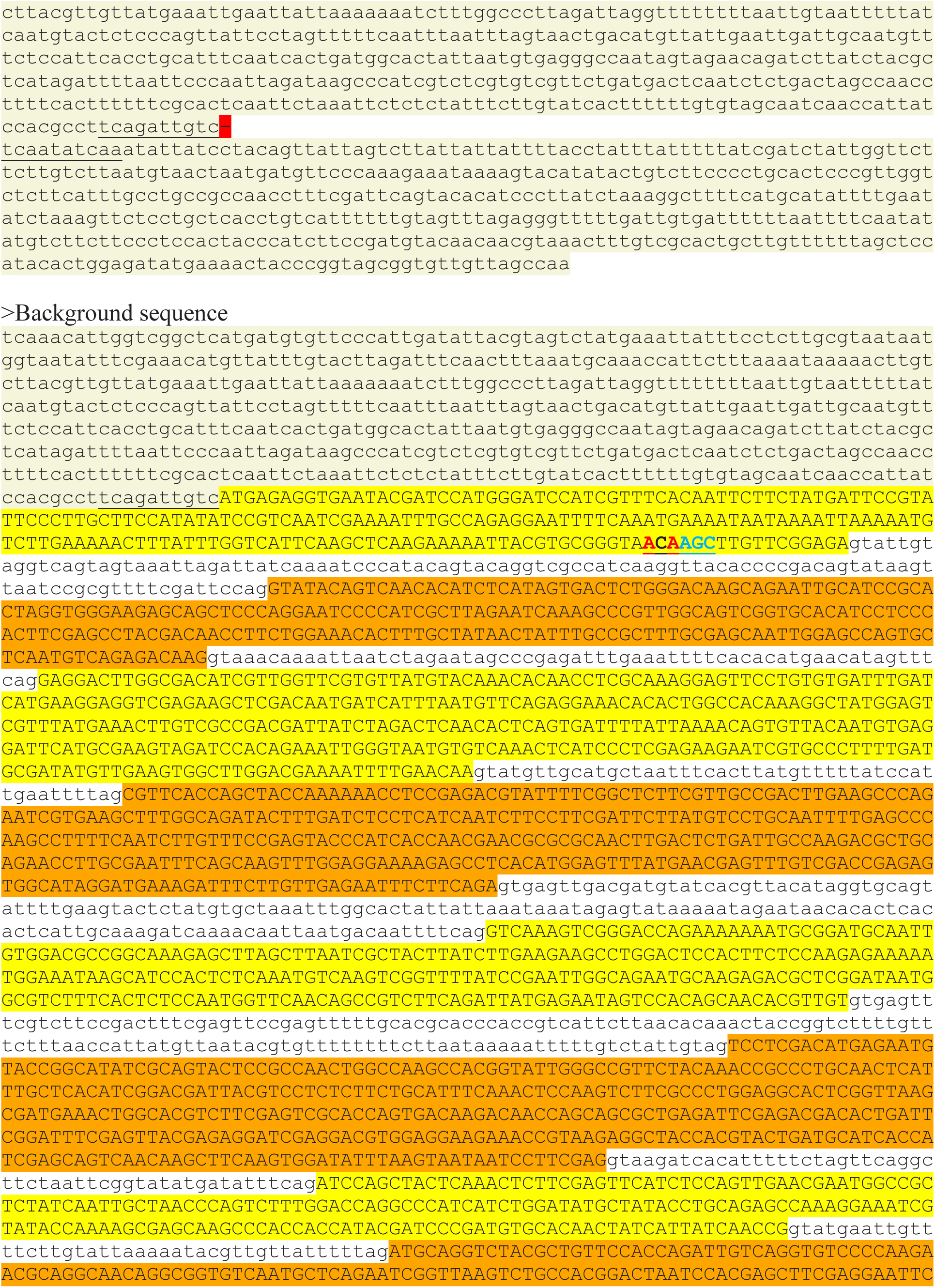
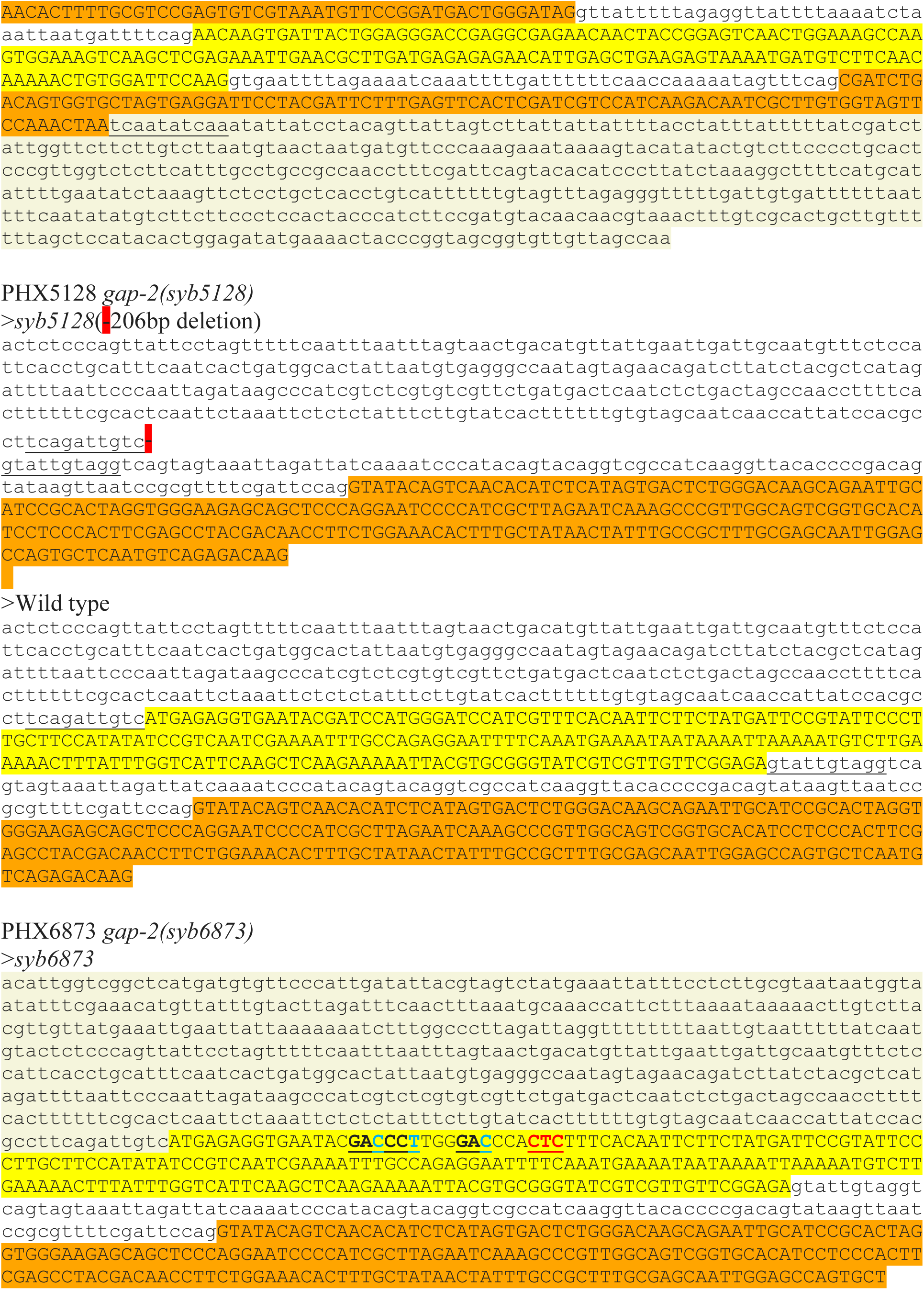
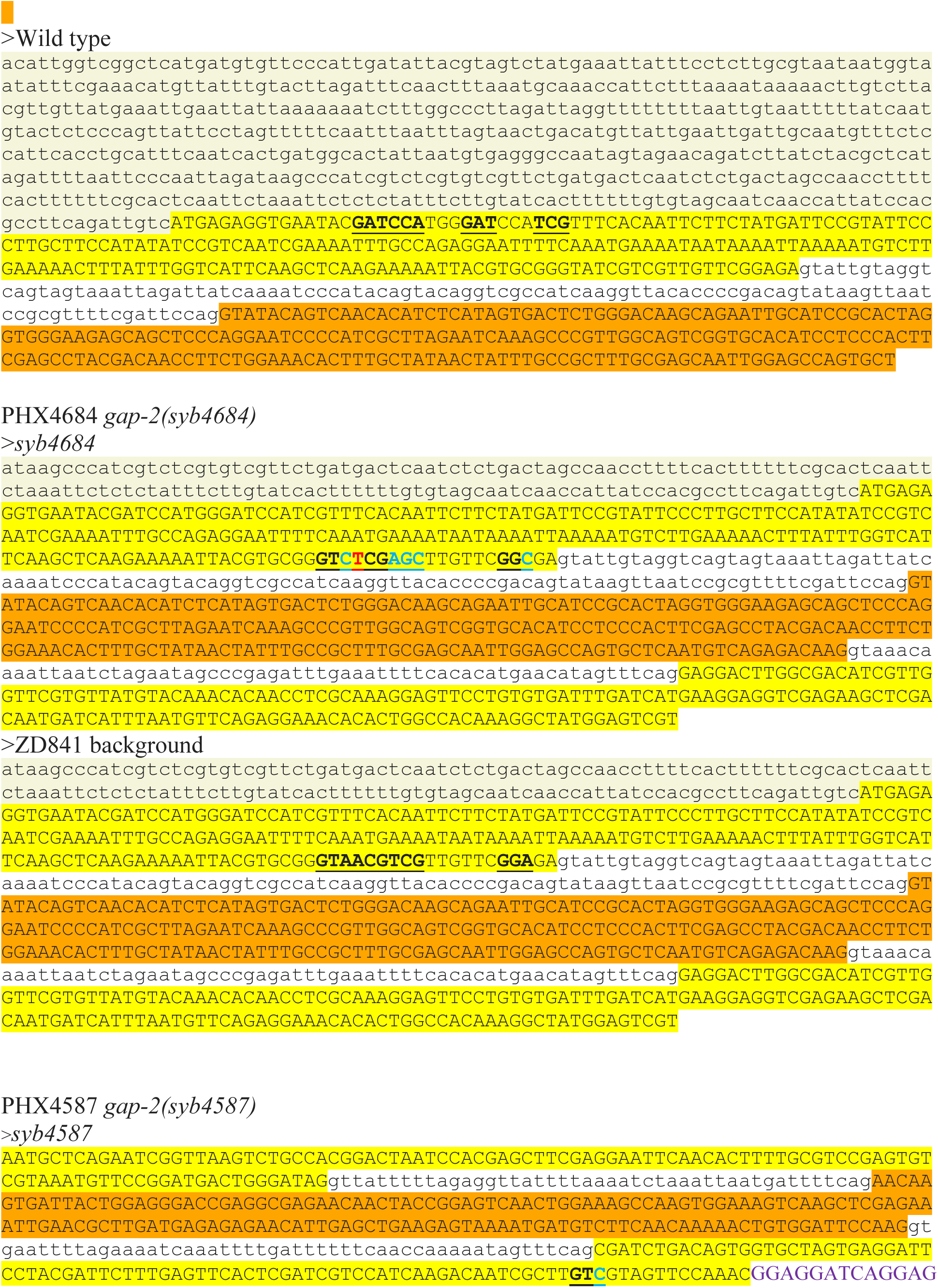
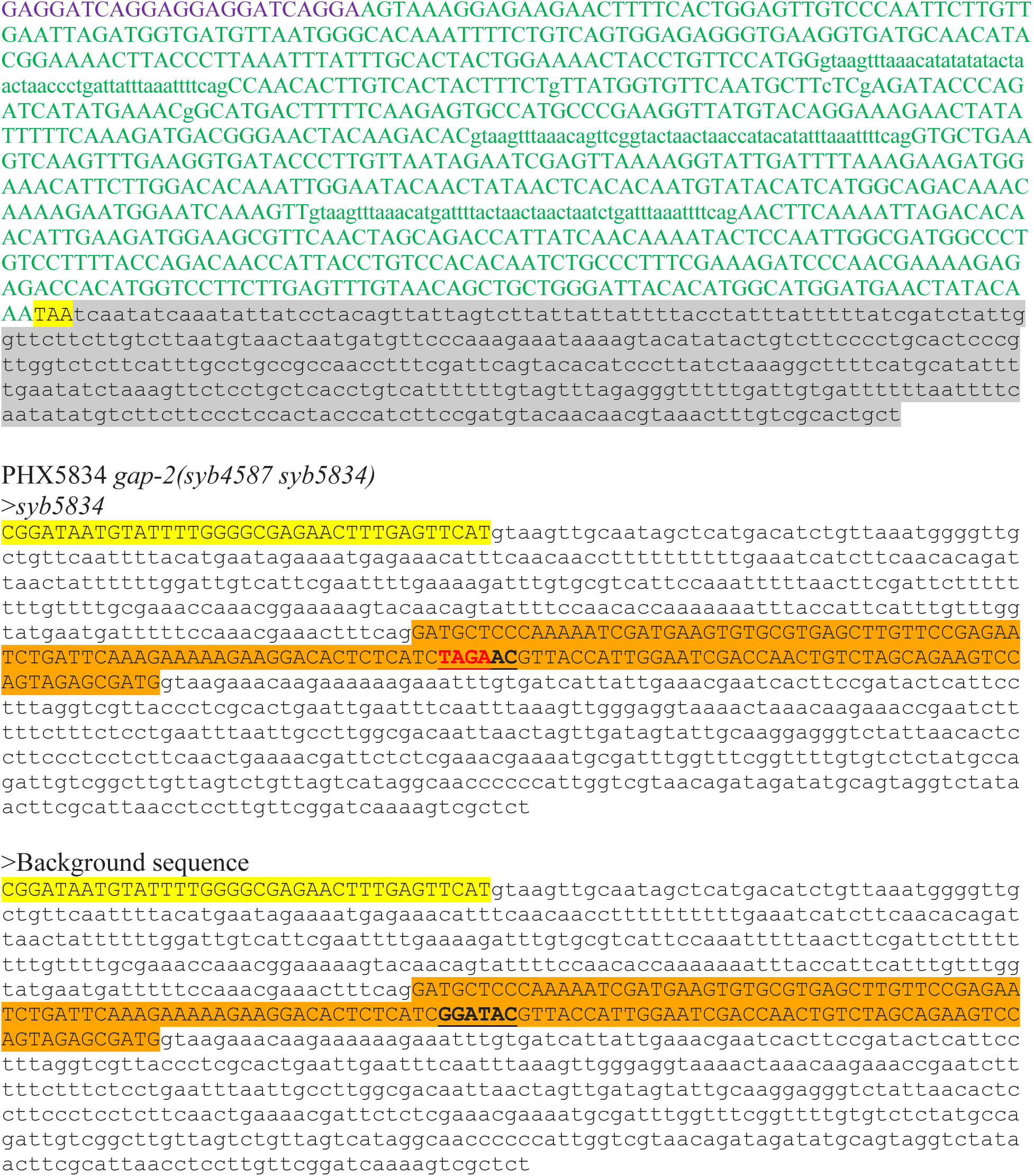

